# Phylogeny of *Paullinia* L. (Paullinieae: Sapindaceae), a diverse genus of lianas with rapid fruit evolution

**DOI:** 10.1101/673988

**Authors:** Joyce G. Chery, Pedro Acevedo-Rodríguez, Carl J Rothfels, Chelsea D. Specht

## Abstract

*Paullinia* L. is a genus of c. 220 mostly Neotropical forest-dwelling lianas that displays a wide diversity of fruit morphologies. *Paullinia* resembles other members of the Paullinieae in being a climber with stipulate compound leaves and paired inflorescence tendrils. However, it is distinct in having capsular fruits with woody, coriaceous, or crustaceous pericarps. While consistent in this basic plan, the pericarps of *Paullinia* fruits are otherwise highly variable—in some species they are winged, whereas in others they are without wings or covered with spines. With the exception of the water-dispersed indehiscent spiny fruits of some members of *Paullinia* sect. *Castanella*, all species are dehiscent, opening their capsules while they are still attached to the branch, to reveal arillate animal-dispersed seeds. Here we present a molecular phylogeny of *Paullinia* derived from 11 molecular markers, including nine novel single-copy nuclear markers amplified by microfluidics PCR. This is the first broadly sampled molecular phylogeny for the genus. *Paullinia* is supported as monophyletic and is sister to *Cardiospermum* L., which together are sister to *Serjania* Mill + *Urvillea* Kunth. We apply this novel phylogenetic hypothesis to test previous infrageneric classifications and to infer that unwinged fruits represent the ancestral condition, from which there were repeated evolutionary transitions and reversals. However, because the seeds of both winged and unwinged fruits are all dispersed by animals, we conclude that the repeated transitions in fruit morphology may relate to visual display strategies to attract animal dispersers, and do not represent transitions to wind dispersal.

## 1. Introduction

*Paullinia* L. (Paullinieae: Sapindaceae) is a genus of c. 220 lianas native to the Neotropics, with one species ranging from tropical sub-saharan Africa to Zimbabwe (Radlkofer, 1933; Irvine, 1961; Medeiros et al., 2016; Acevedo-Rodríguez and Somner 2018). The Amazon region is the center of diversity of the genus (Medeiros et al., 2016) and contains 44% of the described species. Members of *Paullinia* can be identified based on their habit (lianas or vines—seldom erect shrubs), their alternate compound leaves with a terminal leaflet, and their septifragal capsular fruits enclosing arillate seeds. Within this basic morphology there is great variation, particularly in fruit morphology (Radlkofer, 1933, 1895; Acevedo-Rodríguez et al., 2017), degree of leaf dissection, and presence or absence of stems with cambial variants (Bastos et al., 2016; Cunha Neto et al., 2018; Pellissari et al. 2018).

In addition to its taxonomic diversity, morphological disparity, and ecological significance as a prominent component of Neotropical forests (Gentry, 1991), the genus has an extensive history of human utilization. Almost 20% of *Paullinia* species are reported to have ethnobotanical uses by the indigenous peoples of Central and South America, primarily as fish poisons, medicines, and caffeine-rich stimulants (Beck, 1990), and stem cross-sections are used in Brazilian marquetry (Tamaio, 2011). The greatest economic impact, however, is from the caffeine-rich seeds of *P*. *cupana* Kunth, known colloquially as *guaraná*, which is an important international export commodity for Brazil (Erickson et al., 1984).

What we now recognize as *Paullinia* was introduced by Plumier (1693) as “*Clematis”*, and subsequently renamed as “*Cururu*” (Plumier, 1703). Although Plumier (1703) recognized *Paullinia* (*Cururu*) to be distinct from the closely related *Serjania*, Linneaus (1753) included members of *Serjania* within his concept of *Paullinia*. Miller (1754) recognized *Serjania* Mill. as separate from *Paullinia*, and this treatment was further supported by Schumacher (1794), who called for the recognition of fruit morphology as an “essential character”; *Serjania* is easily distinguished from *Paullinia* by its samaroid fruits (rather than capsules). *Paullinia* and *Serjania* were placed in section Paullinieae along with *Urvillea* Kunth and *Cardiospermum* L. (Kunth 1821); this section was later transferred to the rank of tribe by de Candolle (1824) a treatment followed by Radlkofer (1890).

More than a century after *Paullinia* was published, Radlkofer (1895, 1933) reviewed all 91 published names and described a total of 148 species. He subdivided the genus into 13 sections based heavily on fruit characters (Table 1, Figure 1), with the first couplet of his *Conspectus Sectionum* (Radlkofer, 1895, 1933) dividing the genus according to whether the capsules are alate (i.e., winged) or exalate (i.e., without wings; Figure 1 and 2) Finer-scale divisions relied on pericarp morphology and anatomy, inflorescence groupings, number and degree of connation of sepals, and presence of mucilage in leaves (Radlkofer, 1895, 1933). A century later, however, Beck, in his dissertation (1991), concluded that Radlkofer’s (1895, 1933) system was unstable and lacking clear structure. He proposed, instead, that *Paullinia* sensu stricto be reduced to 60 spp., and placed the remaining species into five smaller genera (Beck, 1991). His generic system (Table 1) begins by separating taxa based on seed attachment, then by pericarp morphology and venation. The two large genera, *Paullinia* and *Prancea*, were organized into sections based on pericarp wing and aril characters, respectively. However, as Beck’s (1991) dissertation was never published, Radlkofer’s (1895, 1933) infrageneric classification remains in effect today.

**Table 1.**
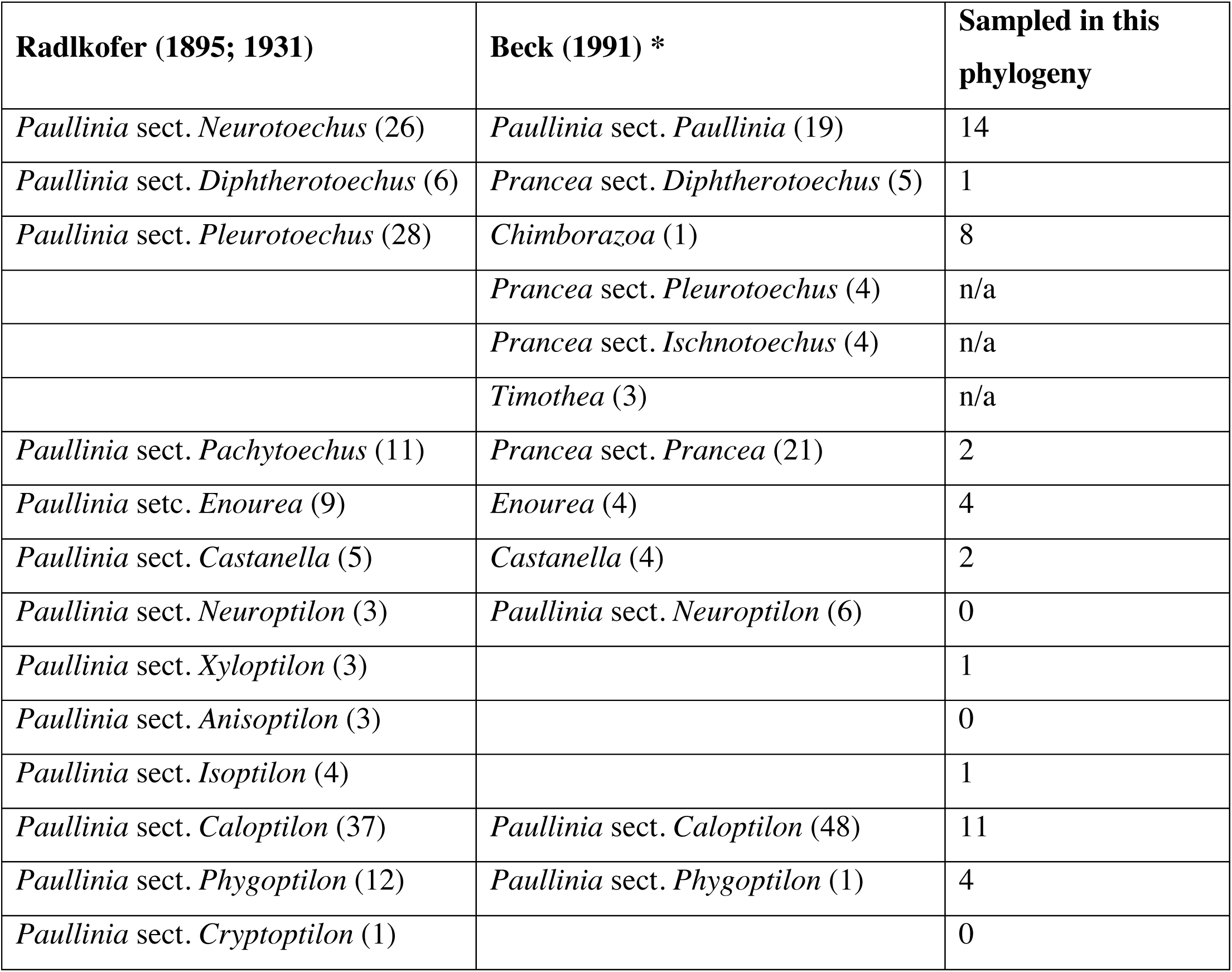
Comparison of the two major classification schemes for *Paullinia*. The number of recognized species are in parentheses. * Beck (1991) classification of *Paullinia* s.s. plus five smaller genera: *Enourea, Castanella, Prancea, Timothea* and *Chimborazoa*.

**Figure 1.**
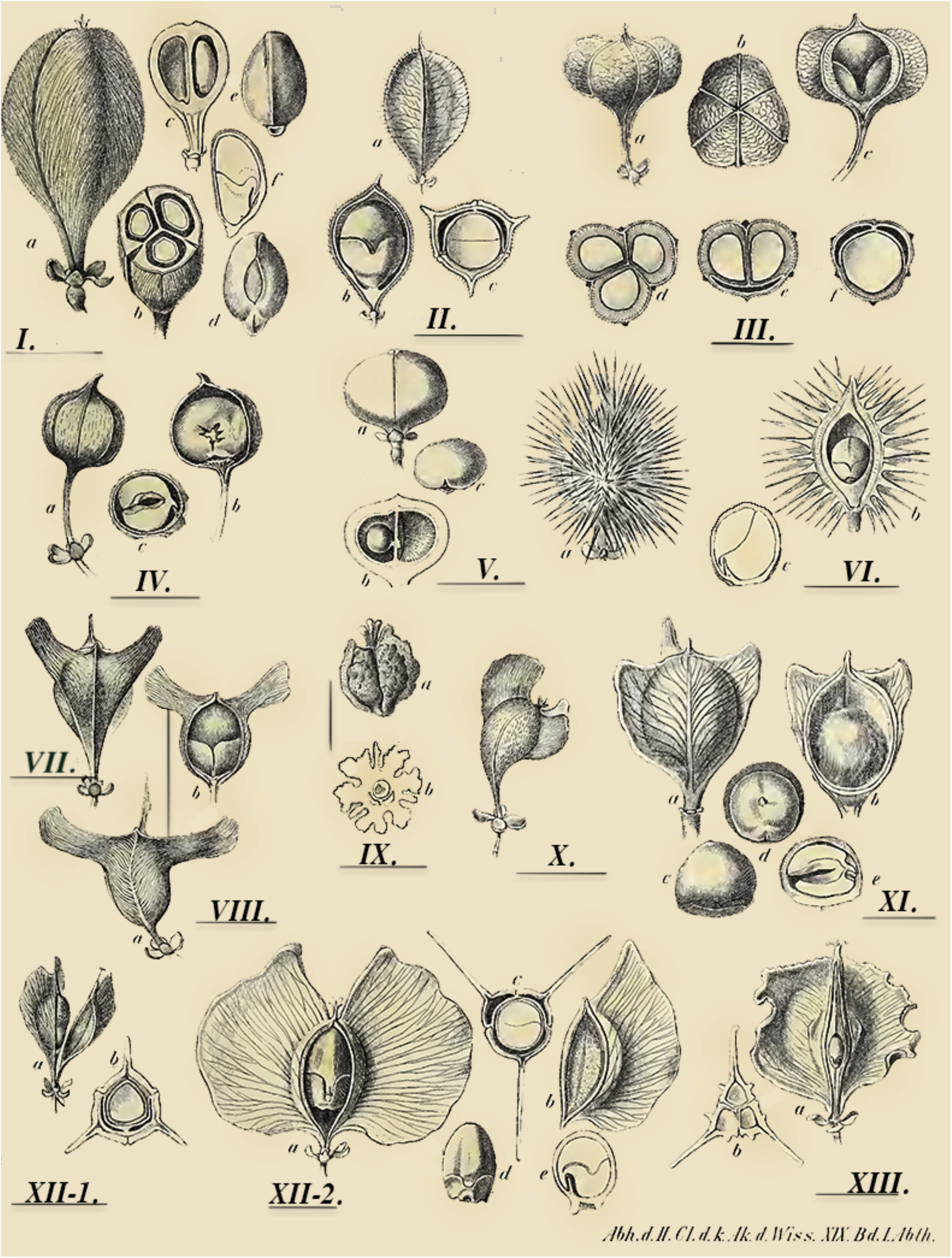
Diversity in fruit morphologies in *Paullinia*; plate modified from Radlkofer (1895) representing his classification system. I. Section *Neurotoechus*, II. Section *Diphtherotoechus*, III. Section *Pleurotoechus*, IV. Section *Pachytoechus*, V. Section *Enourea*, VI. Section *Castanella*, IX. Section *Cryptoptilon*, VIII. Section *Neuroptilon*, VII. Section *Xyloptilon*, X. Section *Anisoptilon*, XI. Section *Isoptilon*, XII. Section *Caloptilon*, XIII. Section *Phygoptilon*.

**Figure 2.**
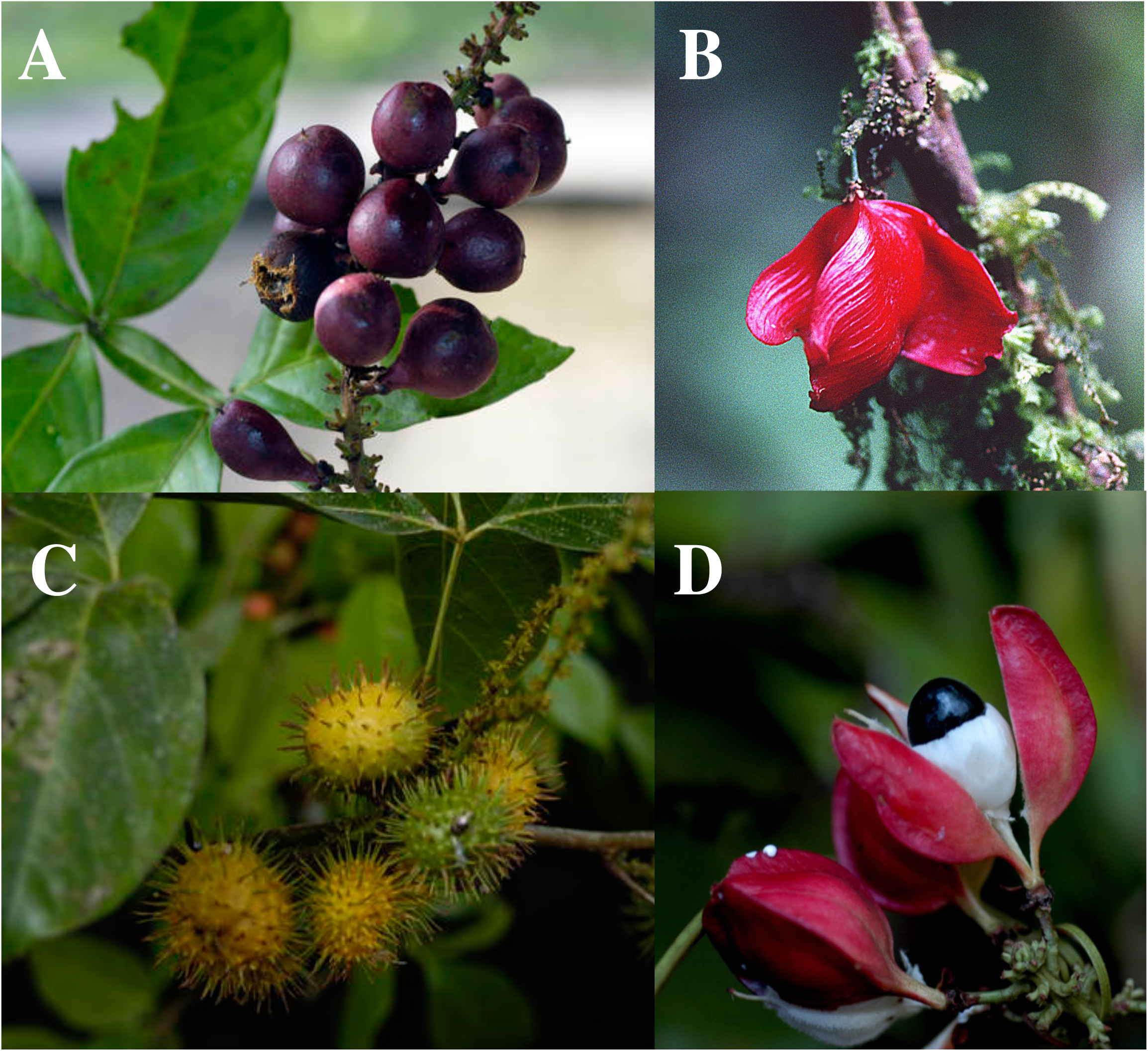
Diversity in fruit morphologies in *Paullinia*. **(A)** dehiscent exalate fruits of *P*. *imberbis* Acevedo 14849, **(B)** dehiscent alate fruits of *P*. *serjaniaefolia* Acevedo 7398, **(C)** indehiscent echinate fruits of *P*. *paullinoides* Acevedo 14860, **(D)** open capsule of the dehiscent alate fruits of *P*. *caloptera* Acevedo 1431, capsule open along septa to reveal black glossy seed covered by a white fleshy aril. All photos by Pedro Acevedo-Rodriguez from collections.nmnh.si.edu.

Until the advent of molecular systematics, which facilitated not only a broad sampling of taxa across the Sapindaceae but allowed testing of proposed affinities based on morphological characters, the monophyly of Paullinieae and the relationships among its genera remained untested. The earliest molecular-based phylogenetic analysis including Sapindaceae (Gadek et al., 1996) did not sample Paullinieae species, but later analyses supported the monophyly of the tribe (Harrington et al., 2005; Buerki et al., 2009; Buerki et al., 2010) and recovered similar relationships to those inferred from cladistic analyses of vegetative and reproductive characters (Acevedo-Rodríguez, 1993) and a hierarchical analysis of wood anatomy using Ward’s clustering algorithm (Klaassen, 1999). Recently, Acevedo-Rodríguez et al. (2017) confirmed the monophyly of Paullinieae (comprising *Thinouia, Lophostigma, Urvillea, Cardiospermum, Serjania*, and *Paullinia*) as one of the four successively nested clades within the greater supertribe Paulliniodae. Paullinieae members are united by their climbing habit, stipulate leaves, and paired inflorescence tendrils (de Candolle, 1824; Radlkofer, 1890; Acevedo-Rodríguez et al., 2017), and can be divided into the “*Paullinia* group” (*Urvillea, Cardiospermum, Paullinia*) and the “*Serjania* group” (*Serjania, Lophostigma* and *Thinouia*), based on whether their fruit are capsules or samaroid schizocarps (Acevedo-Rodríguez 1993); these groups, however, are not monophyletic (Acevedo-Rodríguez et al., 2017).

The history of taxonomic confusion surrounding *Paullinia* and its relatives reveals a lack of macroevolutionary understanding of this lineage, and the reliance on fruit characters highlights the long-standing interest in fruit evolution in the group. However, despite these charismatic features, and the economic and ecological importance of the genus, we still lack a robust phylogenetic framework for *Paullinia*. In this study, we aim to 1) infer the first broadly sampled molecular phylogeny of *Paullinia*; 2) explore taxonomic implications of this phylogeny, and; 3) infer the patterns of fruit evolution within the genus. Specifically, the diversity of fruit morphologies— dehiscent winged and unwinged capsules, and indehiscent fruits with echinate pericarps—are hypothesized to be adaptive features aiding in dispersal (Acevedo-Rodríguez et al., 2017); this *Paullinia* phylogeny allows the reconstruction of transitions among these fruit morphologies and an exploration of their dispersal implications in this species-rich Neotropical genus.

## 2. Materials and Methods

### 2.1 Molecular Marker Selection and Primer Design

Primers for single-copy nuclear markers were designed using the Chery et al. (2017) bioinformatic pipeline, selecting for introns (estimated 500–1100bp) and exons (estimated 400-500bp) by utilizing publicly available Sapindales genomic resources: the *Citrus sinensis* v1.0 genome and the *Dimocarpus longan* transcriptome (Lai and Lin, 2013) (scripts available on github.com/joycecheryPaullinia_Phylogeny). Primers were designed in Primer3 (Koressaar and Remm, 2007; Untergasser et al. 2012) with an optimal Tm of 57°C using the SantaLucia (1998) calculator. These primer pairs were modified by addition of conserved sequence tags for amplification on Fluidigm Access Array (Fluidigm San Francisco, California, USA). Chloroplast primers were taken from Demesure et al. (1995), Fazekas et al. (2008), and Taberlet et al. (1991). In total, 87 primer pairs were tested, targeting a total of 50 single-copy nuclear exons, 24 single-copy nuclear introns (including nine from Chery et al. 2017), and 13 previously published chloroplast markers. Primers were ordered through Eurofins MWG Operon, LLC (Huntsville Alabama, USA). Two modifications to the Fluidigm protocol (Fluidigm PN 100-3770 J1, San Francisco, California, USA) were carried out to enhance amplification success: annealing temperature of 57°C rather than 60°C and use of Phire Hot Start II DNA Polymerase reagents (ThermoFisher Scientific, Pittsburgh, Pennsylvania, USA) rather than the FastStart™ High Fidelity PCR System, dNTPack (Millipore Sigma, St. Louis, Missouri, USA).

### 2.2 Sampling Scheme, DNA Extraction

Taxon sampling prioritized testing Radlkofer’s classification (1895, 1933) and spanning the morphological variation in *Paullinia*, while taking advantage of available silica-dried leaf material, which yield the highest quality DNA extractions. Genomic DNA was extracted from 191 samples (43 herbarium vouchers and 148 silica-dried leaves). CTAB DNA extractions were performed by an Autogen 965 at the Smithsonian Institution Support Center. Extraction quantity was measured by Qubit™ dsDNA HS Assay Kit (ThermoFisher Scientific, Waltham, Massachusetts, USA) at the UC Berkeley DNA sequencing facility. All extractions were diluted to a maximum of 50ng/ul as recommended by the iBEST Genome Resource Core (Moscow, Idaho). Extraction quality of all samples was tested by PCR amplification of ITS, which was sanger-sequenced on an Applied Biosystems 3730xl DNA analyzer at the UC Berkeley DNA Sequencing Facility. Sanger-sequenced reads were trimmed and cleaned in Geneious v.8.0.5 (Biomatters Ltd., Auckland, New Zealand) and aligned using MAFFT local alignment (Katoh and Standley, 2013).

### 2.3 Primer Validation

Primer pairs were tested for amplification in four accessions (*Paullinia turbacensis* Chery 13, *Paullinia* sp. Breedlove 72699, *Paullinia carpopodea* Pace 317, and *Paullinia hystrix* Acevedo 14408) that span the phylogenetic breadth of the genus according to a preliminary ITS phylogeny. Thirty loci that successfully amplified under identical PCR conditions were further pursued. PCR products for these loci were sequenced directly on an ABI 3730x at the Evolutionary Genomics Laboratory at UC Berkeley. Primers were manually removed from sequences and the cleaned reads were aligned using MAFFT (Katoh and Standley, 2013) implemented in Geneious v.8.0.5 (Biomatters Ltd., Auckland, New Zealand) to test for sequence variation among taxa. Two of the thirty loci generated two PCR products and were not pursued further. The final set of target loci included four chloroplast markers (Demesure et al., 1995; Fazekas et al., 2008; Taberlet et al., 1991), thirteen novel single-copy nuclear exon loci, four novel single-copy nuclear intron markers, and seven intron markers developed by Chery et al. (2017; see Appendix A).

### 2.4 Amplification, Sequencing, and Data Processing

DNA extractions, primers, and PCR reagents were sent to the iBEST Genome Core Facility (Moscow, Idaho) for amplification by microfluidic PCR and Illumina sequencing. Samples were run through a Fluidigm 192.24 chip with the standard protocol except for Phire Hot Start II reagents and annealing TM of 57°C. Amplicons were pooled and gel-purified, then run on a fragment analyzer to verify quality. qPCR was performed to determine the quantity of sequenceable libraries, and these were sequenced on 25% of a MiSeq lane.

Illumina Miseq reads were trimmed of reverse primer sequences and demultiplexed by dbcAmplicon (github.com/msettles/dbcAmplicons) by iBEST and additionally cleaned of forward primers and low quality reads with Trimmomatic v.38 (settings: ILLUMINACLIP:TruSeq3-PE-2.fa (this file was modified to include all primers):2:30:10 LEADING: 3 TRAILING:3 SLIDINGWINDOW: 4:15 MINLEN: 36; Bolger et al., 2014). Clean reads were processed through the Fluidigm2PURC pipeline (Blischak et al., 2018). This pipeline is specifically tailored to process Illumina data generated from amplicons and accounts for PCR error and Illumina sequencing errors to predict the likely haplotype(s) for each accession at each locus. The first step merges cleaned paired-end reads using FLASH 2 (Magoc and Salzberg, 2011). These merged reads are processed through PURC (Rothfels et al. 2017), which iteratively clusters reads with the USEARCH cluster_fast algorithm (Edgar, 2010), and detects chimeras using UCHIME’s USEARCH function (Edgar et al., 2011) to generate sequences of haplotype(s) for each accession at each locus. Finally, a maximum likelihood estimate of the number of haplotypes for each accession at each locus is generated by the *crunch cluster* script (Blischak et al., 2018). The output of the Fluidigm2PURC analysis is a MUSCLE alignment (Edgar, 2004) of all haplotypes for each locus. To validate the repeatability of inferred haplotypes (Rothfels et al., 2017), three different PURC regimes were run (denominated “A”, “B”, and “C”), each with four clustering and chimera-killing iterations and a minimum of 10 reads required for a cluster to be retained at each step (corresponding to ∼10x mean coverage). For example, the regime A clustering criteria were .975, .995, .995, .995, meaning that in the first iteration of this regime sequences must be 97.5% identical in order to be clustered together, and these haplotypes (the consensus sequence of each cluster plus any as-yet unclustered sequences) are then fed into the second iteration requiring 99.5% identity, followed by two additional iterations of 99.5% identity each. The following three regimes were implemented: A=.975 .995 .995 .995; B= .995 .995 .995 .975; C= .995 .995 .995 .995). For each locus, maximum likelihood gene trees were inferred in RAxML V. 7.2.8 (Stamatakis, 2014) from the output alignment and putative contaminants and paralogs were removed according to the following workflow: 1) if an accession had multiple haplotypes that formed a monophyletic group, one of these sequences was selected at random; 2) if a putative lineage-specific duplication led to two clades that did not share precisely the same set of taxa (due to inadequate sequencing coverage or other factors), all accessions involved in the duplication were removed from that alignment and; 3) if two or more accessions had identical sequences in all loci, these were treated as contaminants (one for the other) and removed. All scripts are available at github.com/joycecheryPaullinia_Phylogeny.

### 2.5 Phylogenetic Inference

The final concatenated alignment of 11 loci (nine single-copy nuclear markers, plastid *psb*A-*trn*H, and ITS; for accession list see Appendix B) was analyzed by PartitionFinder2 (Lanfear et al., 2016) implemented in CIPRES (Miller et al., 2010) to select the best-fit partitioning scheme and models of evolution for the data (model=mrbayes, linked branch lengths, BIC model selection metric, search algorithm=all). The input to the PartitionFinder2 analysis was the full alignment with each locus designated as its own data subset. A partitioned Bayesian analysis with two runs each of four chains (one cold, three hot; temp=.02) was performed in MrBayes v.3.2.6, sampling every 1000 generations for 10 million generations (Ronquist et al., 2012). The analysis converged with a standard deviation of split frequencies = .008 and the estimated sample size (ESS) of all parameters exceeded 3000. TreeAnnotator v1.10.4 (Bouckaert et al., 2014) was utilized to generate the maximum clade credibility tree using the post-burnin trees from the combined MrBayes runs (Figure 3, 4, 5).

**Figure 3.**
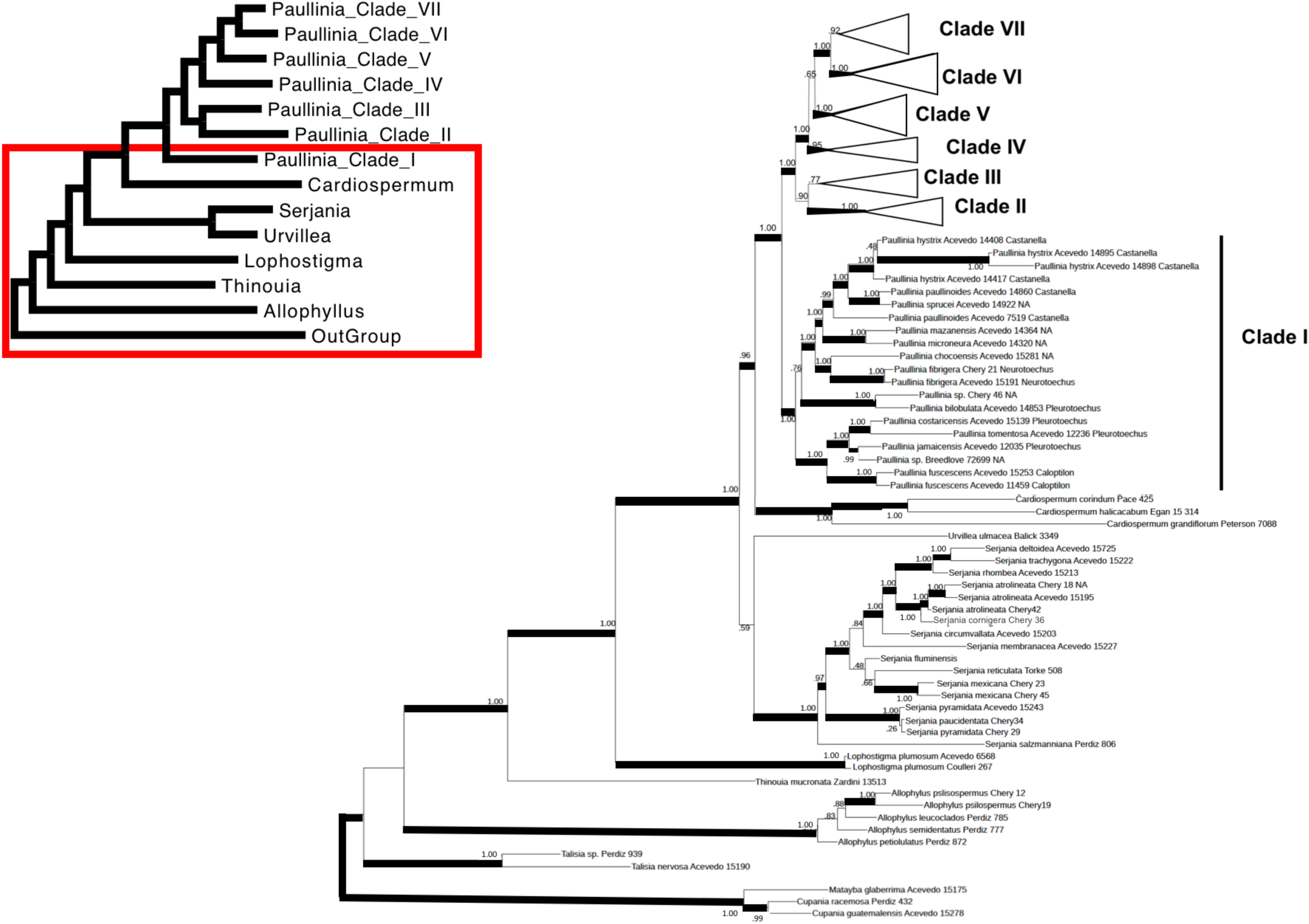
Bayesian maximum clade credibility tree of the outgroup and *Paullinia* Clade I.

**Figure 4.**
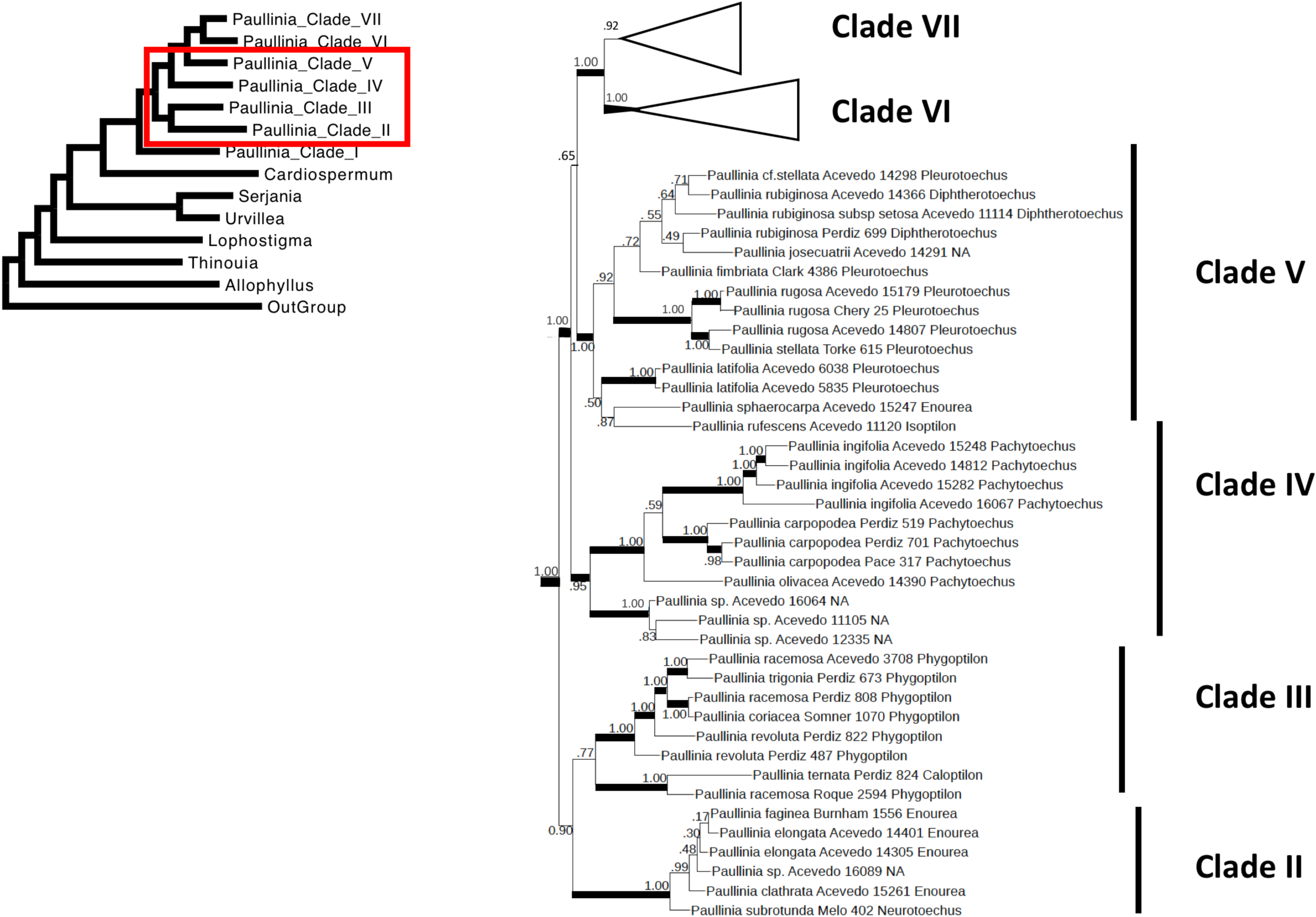
Bayesian maximum clade credibility tree of *Paullinia* Clades II, III, IV and V.

**Figure 5.**
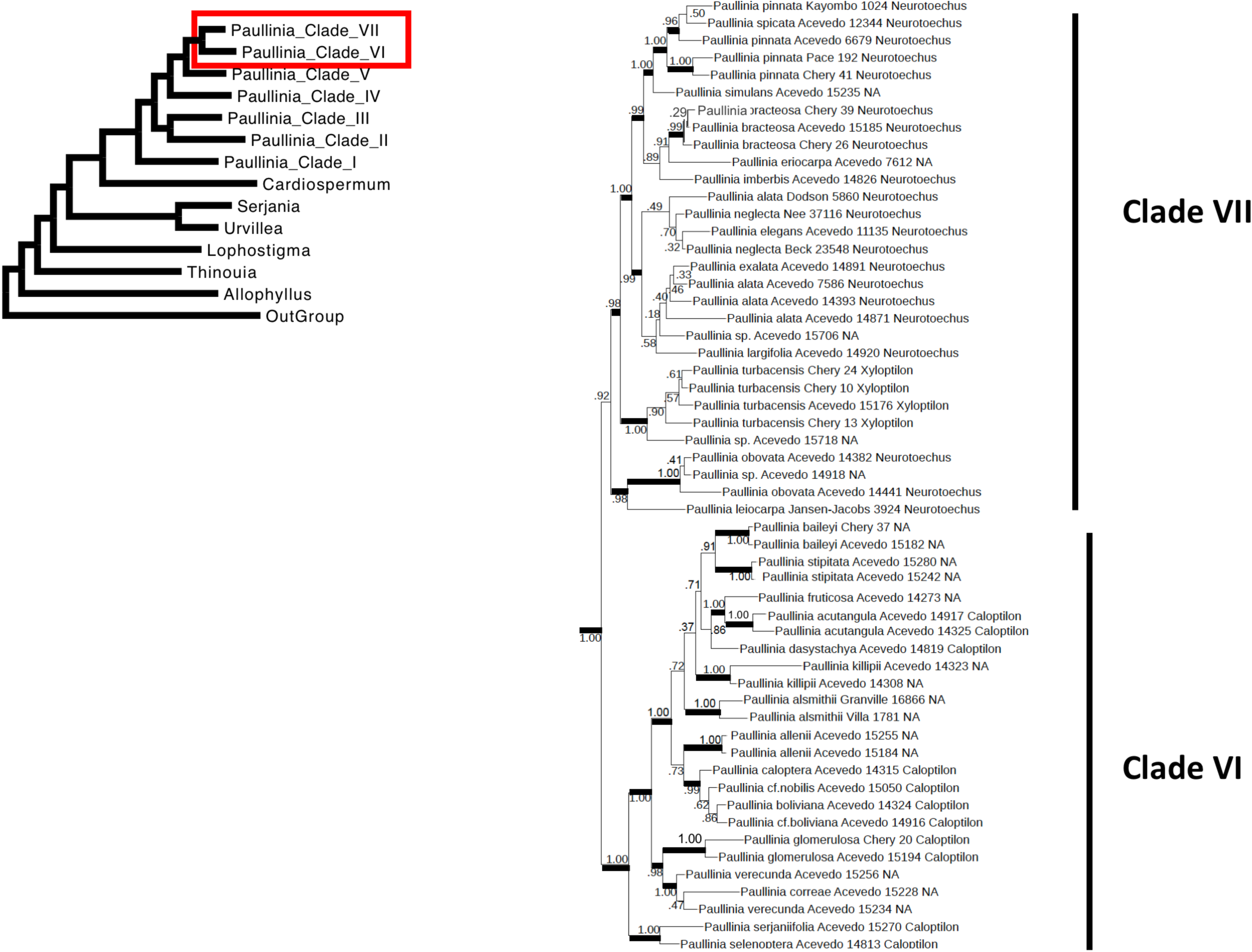
Bayesian maximum clade credibility tree of *Paullinia* Clades VI and VII.

### 2.6 Trait Evolution

*Paullinia* fruit morphologies were categorized as alate (Figure 2B, D), exalate (Figure 2A), or echinate (Figure 2C). Fruit morphology was determined by examining the voucher of each accession in the phylogeny if fruits were present and/or voucher images were available on the Smithsonian Institution Herbarium web database (collections.nmnh.si.edu/search/botany/; accessed 27 February 2019). If fruits were absent from the voucher, but species identity was confirmed by phylogenetic results, other vouchers of that same taxon were evaluated to score fruit morphology for that accession. If the original voucher was sterile, we only sampled a single individual of that taxon, fruit type was recorded from the species descriptions if this is described. The fruit morphology of 12 accessions were unknown and thus these tips were dropped from the phylogeny and excluded in all trait evolution analyses.

To infer patterns of fruit evolution, we estimated ancestral states and the total number of fruit morphology transitions across the phylogeny. Ancestral states and the number of state changes were estimated and visualized by stochastic character mapping using the best-fit model (=symmetric rates; p=.05) along the branches of the Bayesian maximum clade credibility tree using the make.simmap function in the phytools package (Revell, 2012) in R (R Core Team, 2018). Topological and branch length uncertainties were accounted for in the estimation of total average fruit transitions by mapping character histories along the branches of 100 randomly sampled trees from the MrBayes (Ronquist et al., 2012) posterior distribution of trees, and the results were summarized using the describe.simmap function in the phytools package (Revell 2012; R Core Team, 2018). All trees in these analyses were first rendered ultrametric with a relaxed clock model, then pruned down to *Paullinia* tips with fruit morphology data (=102; using the chronos and keep.tip functions, respectively) in the APE package in R (Paradis et al. 2004; R Core Team, 2018). Scripts can be found at github.com/joycechery/Paullinia_Phylogeny and results are in Appendix C.

## 3. Results

### 3.1 Target Selection and Primer validation

DNA extractions ranged from 2.47–50 ug/ul with an average of 26.3 ug/ul (after diluting all extractions to a maximum of 50 ng/ul). Primer validation resulted in 35% success (30 of the 87 primer pairs amplified successfully in the set of test species). The final set of loci selected for microfluidics PCR was 28: four chloroplast, thirteen single-copy nuclear exons, four novel single-copy nuclear intron markers, and seven intron markers published in Chery et al. (2017; see Appendix A).

### 3.2 Sequencing and Data Processing

The Illumina Miseq (25% of a lane) generated 6,500,757 reads spanning 17 of the 28 target loci (the remaining 11 loci did not yield sequence). After additional cleaning with Trimmomatic v.38 (Bolger et al., 2014), 6,279,352 reads remained. Of the 191 samples, 189 produced reads for at least one locus. Only loci with at least 9% success rate across all accessions were pursued. This resulted in 10 loci generated by Miseq and the inclusion of all ITS sequences generated by PCR and Sanger sequencing. The inferred sequences generated by each of the three PURC regimes were consistent (as visualized by maximum likelihood gene trees of all regime haplotypes), suggesting repeatability of the inferred haplotypes across the regimes. PURC regime A typically inferred haplotypes for more accessions so was preferred for nine of the 10 loci. For the remaining locus, regime A generated excessive haplotypes, so regime C was preferred. Two loci were too long for the Miseq paired-end reads to overlap and their alignments contain a central region treated as missing data (“?”).

### 3.3 Phylogenetic Inference

The final concatenated alignment contained 148 OTUs, 814 sequences, 1684 parsimony-informative sites, and 5881 base pairs across nine single-copy nuclear markers (three novel and six from Chery et al. (2017), *psb*A-*trn*H, and ITS. The PartitionFinder2 best scheme favored four partitions as follows: partition one (orange1.1g002083m intron9, orange1.1g027952m intron5, orange1.1g009973m intron5, orange1.1g030977m intron1, orange1.1g036770m intron27): HKY + G; partition two (orange1.1g015495m intron8, orange1.1g016982m intron11, *psb*A-*trn*H): HKY + G; partition three (orange1.1g022777m intron3, orange1.1g019384m intron3); and partition four (ITS): GTR+ G. Gene names are adopted from the *Citrus sinensis* (Rutaceae) v.1.0 genome (Wu et al., 2014) and Chery et al. (2017).

The maximum clade credibility tree (Figure 3, 4, 5) is well resolved with 75% of the nodes having ≥ 95% posterior probability (PP). Rooted with *Cupania* and *Matayba, Allophylus* is sister to the Paullinieae. The first diverging lineage of Paullinieae is *Thinouia*, followed by *Lophostigma*, which is followed by a *Urvillea* + *Serjania* clade that is sister to *Cardiospermum* + *Paullinia*.

Within *Paullinia*, seven clades are described, which roughly correspond to sections sensu Radlkofer (1895, 1931; Figure 3, 4, 5): Clade I (1.0 PP) = sect. *Castanella*, sect. *Caloptilon*, sect. *Pleurotoechus*, and sect. *Neurotoechus*; Clade II (1.0 PP) = sect. *Enourea*; Clade III (0.77 PP) = sect. *Phygoptilon*; Clade IV (0.95 PP) = sect. *Pachytoechus*; Clade V (1.0 PP) = sect. *Pleurotoechus, P*. *rubiginosa* of sect. *Diphtherotoechus, P*. *rufescens* of sect. *Isoptilon*, and *P*. *sphaerocarpa* of sect. *Enourea*; Clade VI (1.0 PP) = sect. *Caloptilon*; and Clade VII (0.92 PP) = sect. *Neurotoechus* and *P*. *turbacensis* of sect. *Xyloptilon*.

### 3.4 Trait Evolution

The ancestral *Paullinia* fruit morphology was reconstructed as exalate (Figure 6), a character that is also found in the sister lineage, *Cardiospermum* s.s.. Seven fruit transitions are inferred on the maximum clade credibility tree: five transitions from exalate to alate (in Clades I, III, IV, VI, and VII), one transition from exalate to echinate at the base of the sect. *Castanella* group in Clade I, and one reversal from alate to exalate in Clade VI. The average number of fruit transitions increases to 8.17 after accounting for topological and branch length uncertainties when character histories of 100 randomly selected trees from the posterior distribution were sampled.

**Figure 6.**
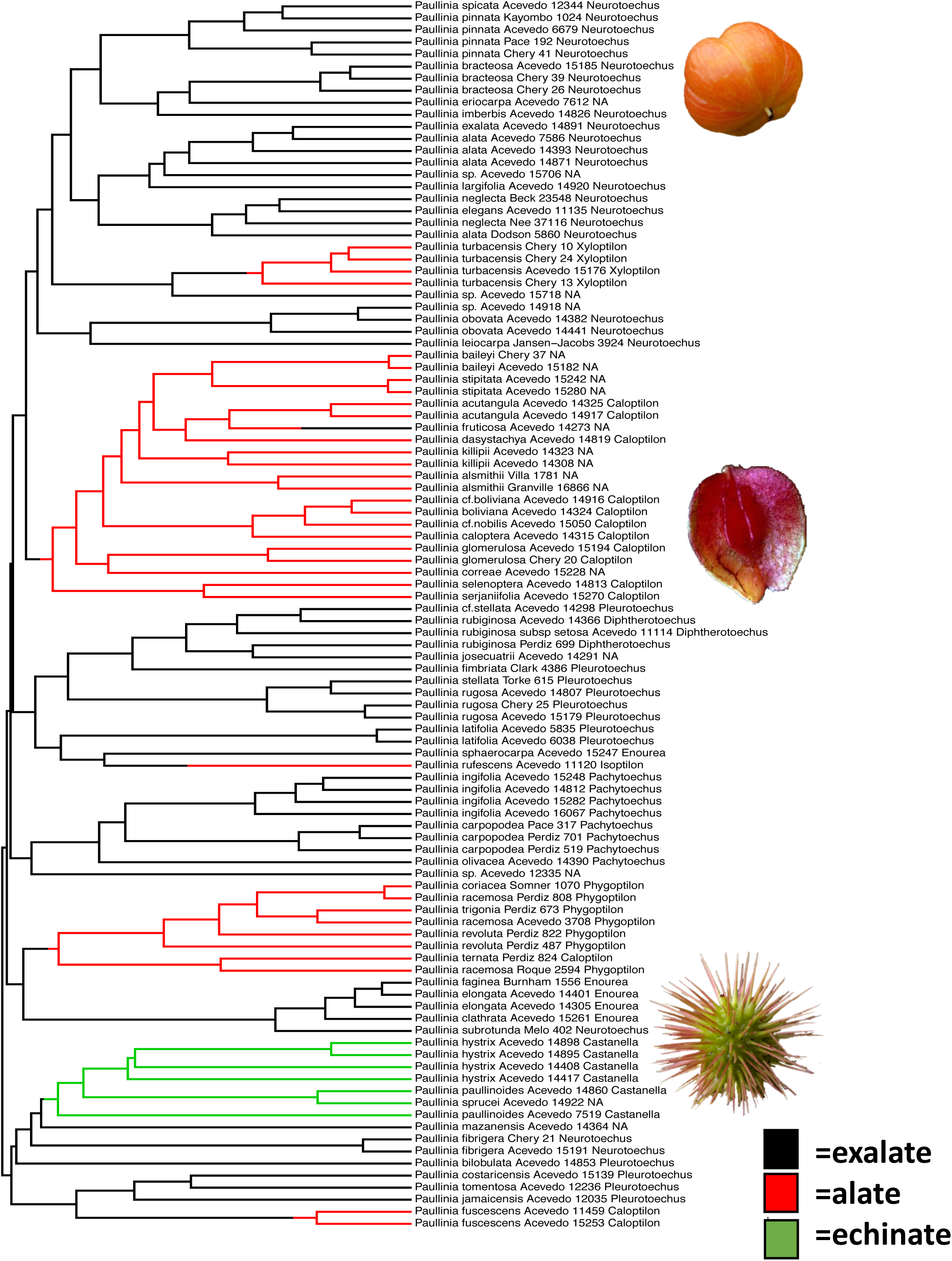
Ancestral state estimations of fruit morphology on the *Paullinia* maximum clade credibility tree.

## 4. Discussion

### 4.1 Taxonomic Implications

The utilization of microfluidics PCR and Illumina sequencing facilitated the generation of the first broadly sampled molecular phylogeny for the genus *Paullinia*. Given the lack of resolution from chloroplast markers for species-level relationships in the genus (Acevedo-Rodríguez et al., 2017; Chery, unpublished), it was necessary to employ rapidly evolving loci in order to generate robust phylogenetic hypotheses to investigate taxonomy and explore morphological evolution. This phylogeny is an improvement from the most recent molecular phylogeny of the Paullinieae, which utilized ITS and the *trnL* intron, where the relationships between *Serjania, Paullinia*, and *Urvillea* were unresolved, and *Cardiospermum* was not monophyletic (Acevedo-Rodríguez et al., 2017). Given the incomplete sampling of this genus, we were unable to test the monophyly of *Cardiospermum*.

The bulk of *Paullinia* sections (sensu Radlkofer 1895, 1933; Table 1) were recovered as monophyletic, thus revealing that many of the morphological characters utilized by classical botanists are synapomorphies for clades and thus carry evolutionary signal as well as being useful for field identification. Some lineages are united by fruit morphology while others are united by various combinations of vegetative characters. Although the analysis presents a great improvement to our macroevolutionary understanding of *Paullina*, increased taxon sampling and/or including more molecular markers is needed to confirm infrageneric relationships and to make new infrageneric circumscriptions, therefore a revised classification will not be proposed until these additional data are incorporated.

The first diverging lineage of *Paullinia*, Clade I, contains sect. *Castanella*, several members of sect. *Pleurotoechus*, and single species from sect. *Neurotoechus* and sect. *Caloptilon*. Clade I is the most diverse in fruit morphology, containing echinate fruits of sect. *Castanella*, exalate fruits of sect. *Neurotoechus* and sect. *Pleurotoechus*, and alate fruits of sect. *Caloptilon*. Here, there is a clear case of a transition in fruit types between closely related species: *P*. *fuscescens* and *P*. *costaricensis* have been described as “difficult to separate vegetatively” (Flora de Nicaragua, accessed on tropicos.org on 11 March 2019), the former with alate fruits and the latter with exalate. All members in this vegetative and reproductively disparate clade display regular stem development (i.e., without cambial variants).

Clade II contains most of sampled sect. *Enourea* species and *P*. *subrotunda* of sect. *Neurotoechus*). Section *Enourea* members have exalate fruits, 5-foliate leaves, and simple stem development (they lack cambial variants). Sister to Clade II, Clade III comprises a monophyletic sect. *Phygoptilon*, except for the inclusion of *P*. *ternata* of sect. *Caloptilon*. Clade III members all have alate fruits and some members have been reported to have successive cambia (Neto et al., 2018). Clade IV consists of a monophyletic sect. *Pachytoechus*, all with exalate fruit.

Clade V comprises half of sampled sect. *Pleurotoechus, P*. *rubiginosa* of sect. *Diphtherotoechus, P*. *rufescens* of sect. *Isoptilon*, and *P*. *josecuatrii*. Fruit morphology varies from exalate and sharply triangular (*P*. *rubiginosa*), to exalate and globose (sect. *Pleurotoechus*), to alate with large wings (*P*. *rufescens*). The group is united by the presence of the phloem wedge cambial variant in most members (Chery, manuscript in prep.).

Clade VI houses most of the sampled sect. *Caloptilon* species. All species in this clade have alate fruits and compound leaves that are 5-foliate and pinnate or highly dissected (e.g., *P*. *glomerulosa*). In some lineages within Clade VI, there are transitions in vegetative characters between closely related species. For example, the 5-foliate and phloem wedge cambial variant species *P*. *baileyi* was recovered as sister to the 3-jugate (or ternate-pinnate *sensu* Croat 1976) and compound-stem species *P*. *stipitata*.

Clade VII houses most of the sampled sect. *Neurotoechus* and the only sampled representative of sect. *Xyloptilon* (*P*. *turbacensis*). This clade is united by the presence of ternate or 5-foliate pinnate compound leaves and compound or phloem wedge cambial variants. Embedded within this group is a clade characterized by the presence of cauliflorous inflorescences (*P*. *alata, P*. *exalata, P*. *largifolia*). All species in Clade VII have pyriform exalate fruits except *P*. *turbacensis* of sect. *Xyloptilon* that has fruits with small wings.

### 4.2 Fruit Evolution

Fruit morphology has been the most important character used to distinguish *Paullinia* from closely related genera and to identity and place taxa within the genus, however the patterns of evolution of *Paullinia* fruit have not been previously explored. The ancestral fruit morphology is inferred to be exalate, and seven transitions are inferred on the maximum clade credibility tree: five transitions from exalate to alate, one transition from exalate to echinate, and one reversal from alate to exalate. The repeated gain of wings might suggest transitions from animal to wind dispersal, which would be consistent with wind-dispersed fruits promoting species diversification (e.g., the evolution of schizocarp fruits with high number of carpels correlated with speciation in the eumalvoids, Malvaceae; Areces-Berazain and Ackerman 2017). This conflicts with the long-standing hypothesis that the radiation of angiosperms is due in part to the evolution of animal-dispersed diaspores that facilitate propagules founding new populations where speciation by isolation can occur (reviewed by Eriksson and Bremer, 1991; but see Herrera, 1989). However, although the differences in fruit morphologies in *Paullinia* superficially imply shifts in dispersal syndromes, field observations reveal that all *Paullinia* fruits open and dehisce their seeds while still attached to the branch, with the exception of some members of the “riparia group” (sect. *Castanella*), which have fruits that are echinate, indehiscent (in at least one species), and water dispersed (Acevedo-Rodríguez, personal observation). The pericarp of the dehiscent capsules are usually reddish (Weckerle and Rutishauser, 2005), and open to display three black glossy seeds covered by a fleshy white aril for consumption. The conspicuousness of the color contrast of the pericarp-seed-aril complex against the forest canopy attracts bird dispersers (van der Pijl, 1982; Howe and Smallwood, 2003; Schmidt et al., 2004). Experiments with wild-caught and hand-raised birds demonstrated significant preference for red fruits (Duan et al., 2014). This exact color contrast is also exhibited by the ovuliferous cones of the conifer genus *Phyllocladus* (Podocarpaceae), which is also bird dispersed (Contreras et al., 2017). If the alate fruits of *Paullinia* were wind dispersed, we would expect them to not open on the branch, and for there to be no aril, such as in *Serjania*; however, the seeds of all the dehiscent-capsuled species of *Paullinia* are arillate. Other animal dispersers such as rodents are reported for *P*. *sphaerocarpa* Rich. ex Juss. (Weckerle and Rutishauser, 2005). The stability of the color-contrast and arillate seeds syndrome across the dehiscent species in *Paullinia* supports the observation that both alate and exalate fruits are animal dispersed, most probably by birds.

If the dispersal unit is the arillate seed rather than the entire alate capsule, then it follows that the fruit wings may function as an attractant. For example, perhaps the wings function to increase the size of the fruit, providing a more attractive visual display to dispersers. The mutualistic relationship between *Paullinia* and birds is heightened in the dry season when most other plants are not producing fruit (Barro Colorado Island, Panama: Croat, 1978; Leigh, 1999). By reaching peak fruiting at this relatively scarce season, birds are most likely to take advantage of this available food resource (Leigh, 1999), facilitating successful dispersal of seeds.

Water dispersal may also be important in *Paullinia*. At least one species, *P*. *hystrix* of sect. *Castanella*, has been reported to be water dispersed, containing green echinate capsules with septa that are slightly thickened and consist of spongy tissue that allows the seed to float (Weckerle and Rutishauser, 2005). Additionally, *P*. *sprucei* of sect. *Castanella* is also water dispersed (Acevedo-Rodriguez personal observation). Another member of this section, *P*. *paullinioides*, however, has red dehiscent echinate capsules. The dispersal syndrome of other *Castanella* members is unknown, however the presence of yellow to green capsules (Figure 2C) in several members is notable because they do not display the optimal color/color contrast for bird dispersal (Duan et al., 2014). Water dispersal has also been observed in the distantly related *P*. *clathrata* (Weckerle and Rutishauser, 2005).

The repeated transitions in fruit morphology across the phylogeny suggest an advantage associated with dispersal, however in the absence of direct field observations of the preferences of birds for alate vs. exalate fruits, and the dispersal success difference between birds vs. water dispersal, this remains an open question.

## 5. Conclusions

Here we present the first molecular phylogeny of *Paullinia* based on a large molecular dataset, consisting of a set of novel rapidly evolving single-copy nuclear markers and two commonly used markers. *Paullinia* is recovered with high support as monophyletic and sister to *Cardiospermum*, which together are sister to *Serjania* + *Urvillea*. The ancestral condition of fruit morphology in *Paullinia* is reconstructed as exalate, and at least seven fruit morphology transitions are estimated in the genus. Although it is tempting to ascribe the evolutionary trend of gain of wings to a transition from animal to wind dispersal, the arillate seeds of both alate and exalate fruits are dispersed by animals; therefore, we conclude that the repeated transitions in fruit morphology represent different strategies to enhance visual display to attract a diversity of bird dispersers. Future research confirming precise bird preferences of different *Paullinia* fruits could uncover the advantages conferred by diverse fruit morphologies.

## Glossary

Alate: winged
Exalate: without wings
Echinate: spiny protrusions on pericarp layer

## Acknowledgements

Thank you to Morgan Gostel and Daniel New for technical support with the microfluidics PCR run and to Michael May for assistance with the ancestral state estimations. We would also like to acknowledge Gabriel Fox from the Smithsonian Museum Support Center and the Laboratory of Analytical Biology for molecular lab assistance, and those who facilitated J.G.C plant collections; Melissa Cano and Oris Acevedo (Barro Colorado Island, Panama) and Holy Forbes (UC Botanical Gardens).

