## Appendix C for "Phylogeny of *Paullinia* L. (Paullinieae: Sapindaceae), a diverse genus of lianas with rapid fruit evolution"

**Appendix C.** Results of 100 stochastic character mapping analyses:

- describe.simmap(SYMmap)

100 trees with a mapped discrete character with states:

0, 1, 2

trees have 8.28 changes between states on average

changes are of the following types:

0,1	0,2	1,0	1,2	2,0	2,1
x->y	5.71	1.03	1.49	0	0.05	0

mean total time spent in each state is:

0	1	2	total

raw 6.1623660	2.9654566	0.74865807	9.876481

prop 0.6239176	0.3003653	0.07571713	1.000000

- [countSimmap(SYMmap)]

**Tree	Number of transitions	(0,0) (0,1) (0,2) (1,0) (1,1) (1,2) (2,0) (2,1) (2,2)**

[1,] 7	0	5	1	1	0	0	0	0	0

[2,] 8	0	6	1	1	0	0	0	0	0

[3,] 7	0	5	1	1	0	0	0	0	0

[4,] 7	0	5	1	1	0	0	0	0	0

[5,] 8	0	5	2	1	0	0	0	0	0

[6,] 9	0	6	1	2	0	0	0	0	0

[7,] 8	0	6	1	1	0	0	0	0	0

[8,] 7	0	5	1	1	0	0	0	0	0

[9,] 7	0	5	1	1	0	0	0	0	0

[10,] 7	0	5	1	1	0	0	0	0	0

[11,] 10	0	7	1	2	0	0	0	0	0

[12,] 8	0	6	1	1	0	0	0	0	0

[13,] 7	0	5	1	1	0	0	0	0	0

[14,] 8	0	5	1	2	0	0	0	0	0

[15,] 7	0	5	1	1	0	0	0	0	0

[16,] 8	0	6	1	1	0	0	0	0	0

[17,] 7	0	5	1	1	0	0	0	0	0

[18,] 7	0	5	1	1	0	0	0	0	0

[19,] 7	0	5	1	1	0	0	0	0	0

[20,] 8	0	5	2	1	0	0	0	0	0

[21,] 8	0	5	1	2	0	0	0	0	0

[22,] 9	0	7	1	1	0	0	0	0	0

[23,] 8	0	5	1	2	0	0	0	0	0

[24,] 12	0	8	1	3	0	0	0	0	0

[25,] 7	0	5	1	1	0	0	0	0	0

[26,] 7	0	5	1	1	0	0	0	0	0

[27,] 7	0	5	1	1	0	0	0	0	0

[28,] 8	0	5	1	2	0	0	0	0	0

[29,] 7	0	5	1	1	0	0	0	0	0

[30,] 8	0	6	1	1	0	0	0	0	0

[31,] 9	0	6	1	2	0	0	0	0	0

[32,] 7	0	5	1	1	0	0	0	0	0

[33,] 8	0	6	1	1	0	0	0	0	0

[34,] 9	0	6	1	2	0	0	0	0	0

[35,] 8	0	5	1	2	0	0	0	0	0

[36,] 7	0	5	1	1	0	0	0	0	0

[37,] 7	0	5	1	1	0	0	0	0	0

[38,] 7	0	5	1	1	0	0	0	0	0

[39,] 9	0	5	1	3	0	0	0	0	0

[40,] 8	0	6	1	1	0	0	0	0	0

[41,] 7	0	5	1	1	0	0	0	0	0

[42,] 8	0	6	1	1	0	0	0	0	0

[43,] 7	0	5	1	1	0	0	0	0	0

[44,] 7	0	5	1	1	0	0	0	0	0

[45,] 8	0	6	1	1	0	0	0	0	0

[46,] 8	0	5	1	2	0	0	0	0	0

[47,] 9	0	6	1	2	0	0	0	0	0

[48,] 11	0	6	1	4	0	0	0	0	0

[49,] 9	0	6	1	2	0	0	0	0	0

[50,] 12	0	7	1	4	0	0	0	0	0

[51,] 7	0	5	1	1	0	0	0	0	0

[52,] 10	0	7	1	2	0	0	0	0	0

[53,] 7	0	5	1	1	0	0	0	0	0

[54,] 9	0	7	1	1	0	0	0	0	0

[55,] 8	0	5	1	2	0	0	0	0	0

[56,] 7	0	5	1	1	0	0	0	0	0

[57,] 8	0	6	1	1	0	0	0	0	0

[58,] 8	0	5	1	1	0	0	1	0	0

[59,] 8	0	6	1	1	0	0	0	0	0

[60,] 11	0	8	1	2	0	0	0	0	0

[61,] 9	0	6	1	1	0	0	1	0	0

[62,] 8	0	6	1	1	0	0	0	0	0

[63,] 10	0	6	1	3	0	0	0	0	0

[64,] 11	0	7	1	2	0	0	1	0	0

[65,] 9	0	6	1	2	0	0	0	0	0

[66,] 8	0	6	1	1	0	0	0	0	0

[67,] 9	0	6	1	2	0	0	0	0	0

[68,] 7	0	5	1	1	0	0	0	0	0

[69,] 8	0	5	1	1	0	0	1	0	0

[70,] 11	0	8	1	2	0	0	0	0	0

[71,] 10	0	7	1	2	0	0	0	0	0

[72,] 7	0	5	1	1	0	0	0	0	0

[73,] 7	0	5	1	1	0	0	0	0	0

[74,] 9	0	6	1	2	0	0	0	0	0

[75,] 9	0	6	1	2	0	0	0	0	0

[76,] 8	0	5	1	2	0	0	0	0	0

[77,] 10	0	7	1	2	0	0	0	0	0

[78,] 8	0	5	1	2	0	0	0	0	0

[79,] 10	0	5	1	4	0	0	0	0	0

[80,] 11	0	7	1	3	0	0	0	0	0

[81,] 8	0	6	1	1	0	0	0	0	0

[82,] 9	0	7	1	1	0	0	0	0	0

[83,] 9	0	7	1	1	0	0	0	0	0

[84,] 12	0	7	1	4	0	0	0	0	0

[85,] 8	0	6	1	1	0	0	0	0	0

[86,] 8	0	5	2	1	0	0	0	0	0

[87,] 7	0	5	1	1	0	0	0	0	0

[88,] 10	0	7	1	2	0	0	0	0	0

[89,] 10	0	7	1	2	0	0	0	0	0

[90,] 8	0	6	1	1	0	0	0	0	0

[91,] 10	0	7	1	2	0	0	0	0	0

[92,] 7	0	5	1	1	0	0	0	0	0

[93,] 8	0	6	1	1	0	0	0	0	0

[94,] 7	0	5	1	1	0	0	0	0	0

[95,] 7	0	5	1	1	0	0	0	0	0

[96,] 7	0	5	1	1	0	0	0	0	0

[97,] 7	0	5	1	1	0	0	0	0	0

[98,] 10	0	7	1	2	0	0	0	0	0

[99,] 8	0	5	1	1	0	0	1	0	0

[100,] 8	0	6	1	1	0	0	0	0	0
