## Appendix B for "Phylogeny of *Paullinia* L. (Paullinieae: Sapindaceae), a diverse genus of lianas with rapid fruit evolution"

**Appendix B:** List of accessions

| ***Taxon*** |  | **Collector** | **Collection No.** | **Herbarium** | **Herbarium Catalog No.** | **Country** |
| --- | --- | --- | --- | --- | --- | --- |
| *Allophylus psilsospermus* | Radlk. | Chery | 19 | PMA | - | Panama |
| *Allophylus leucoclados* | Radlk. | Perdiz | 785 | US | 3627807 | Brazil |
| *Allophylus petiolulatus* | Radlk. | Perdiz | 872 | US | 3627806 | Brazil |
| *Allophylus psilsospermus* | Radlk. | Chery | 12 | PMA | - | Panama |
| *Allophylus semidentatus* | (Miq.) Radlk. | Perdiz | 777 | US | 3627813 | Brazil |
| *Cardiospermum corindum* | L. | Pace | 425 | US | 3677380 | Mexico |
| *Cardiospermum grandiflorum* | Sw. | Peterson | 7088 | US | 3594402 | Panama |
| *Cardiospermum halicacabum* | L. | Egan | 15-314 | US | 3678585 | United States |
| *Cupania guatemalensis* | Radlk. | Acevedo | 15278 | US | 3668963 | Panama |
| *Cupania racemosa* | Radlk. | Perdiz | 432 | US | 3627812 | Brazil |
| *Lophostigma plumosum* | Radlk. | Coulleri | 267 | US | 3628138 | Bolivia |
| *Lophostigma plumosum* | Radlk. | Acevedo | 6568 | US | 3295140 | Bolivia |
| *Matayba glaberrima* | Radlk. | Acevedo | 15175 | US | 3691476 | Panama |
| *Paullinia acutangula* | (Ruiz & Pavón) Pers. | Acevedo | 14325 | US | 3635026 | Peru |
| *Paullinia acutangula* | (Ruiz & Pavón) Pers. | Acevedo | 14917 | US | - | Brazil |
| *Paullinia alata* | (Ruiz & Pavón) G. Don | Acevedo | 7586 | US | 3330090 | Ecuador |
| *Paullinia alata* | (Ruiz & Pavón) G. Don | Acevedo | 14393 | US | 3620749 | Peru |
| *Paullinia alata* | (Ruiz & Pavón) G. Don | Dodson | 5860 | US | 2843911 | Ecuador |
| *Paullinia allenii* | Standl. | Acevedo | 15184 | US | 3691471 | Panama |
| *Paullinia allenii* | Standl. | Acevedo | 15255 | US | 3683522 | Panama |
| *Paullinia alsmithii* | Macbr. | Villa | 1781 | US | 3472337 | Ecuador |
| *Paullinia alsmithii* | Macbr. | Granville | 16866 | US | 3523305 | French Guiana |
| *Paullinia baileyi* | Standl. | Chery | 37 | PMA | - | Panama |
| *Paullinia baileyi* | Standl. | Acevedo | 15182 | US | 3691470 | Panama |
| *Paullinia bilobulata* | Radlk. | Acevedo | 14853 | US | - | Brazil |
| *Paullinia boliviana* | Radlk. | Acevedo | 14324 | US | 3620721 | Peru |
| *Paullinia bracetosa* | Radlk. | Chery | 39 | PMA | - | Panama |
| *Paullinia bracteosa* | Radlk. | Acevedo | 15185 | US | 3691469 | Peru |
| *Paullinia bracteosa* | Radlk. | Chery | 26 | PMA |  | Panama |
| *Paullinia caloptera* | Radlk. | Acevedo | 14315 | US | 3635021 | Peru |
| *Paullinia carpopodea* | Camb. | Perdiz | 701 | CEPEC |  | Brazil |
| *Paullinia carpopodea* | Camb. | Perdiz | 519 | US | 3627811 | Brazil |
| *Paullinia carpopodea* | Camb. | Pace | 317 | US |  |  |
| *Paullinia cf. alata* | (Ruiz & Pavón) G. Don | Acevedo | 14871 | US | 3680205 | Brazil |
| *Paullinia cf. boliviana* | Radlk. | Acevedo | 14916 | US | - | Brazil |
| *Paullinia cf. stellata* | Radlk. | Acevedo | 14298 | US | 3629599 | Peru |
| *Paullinia cf.nobilis* | Radlk. | Acevedo | 15050 | US | 3680336 | Brazil |
| *Paullinia chocoensis* | Cuatre. | Acevedo | 15281 | US | 3668966 | Panama |
| *Paullinia clathrata* | Radlk. | Acevedo | 15261 | US | 3683515 | Panama |
| *Paullinia coriacea* | Casar. | Somner | 1070 | RBR | 30803 | Brazil |
| *Paullinia correae* | Croat | Acevedo | 15228 | US | 3691477 | Panama |
| *Paullinia costaricensis* | Radlk. | Acevedo | 15139 | US | 3582595 | Mexico |
| *Paullinia dasystachya* | Radlk. | Acevedo | 14819 | US | 3680192 | Brazil |
| *Paullinia elegans* | Cambess. | Acevedo | 11135 | US | - | Bolivia |
| *Paullinia elongata* | Radlk. | Acevedo | 14305 | US | 3625010 | Peru |
| *Paullinia elongata* | Radlk. | Acevedo | 14401 | US | 3620757 | Peru |
| *Paullinia eriocarpa* | Tria. & Planch. | Acevedo | 7612 | US | 3330104 | Ecuador |
| *Paullinia exalata* | Radlk. | Acevedo | 14891 | US | 3680199 | Brazil |
| *Paullinia faginea* | (Tria. & Planch.) Radlk. | Burnham | 1556 | US | 3381558 | Ecuador |
| *Paullinia fibrigera* | Radlk. | Chery | 21 | PMA | - | Panama |
| *Paullinia fibrigera* | Radlk. | Acevedo | 15191 | US | 3668964 | Panama |
| *Paullinia fimbriata* | Radlk. | Clark | 4386 | US | 3698347 | Ecuador |
| *Paullinia fruticosa* |  | Acevedo | 14273 | US | - | Peru |
| *Paullinia fuscescens* | Kunth | Acevedo | 11459 | US | 3429796 | U.S.Virgin Islands |
| *Paullinia fuscescens* | Kunth | Acevedo | 15253 | US | 3683516 | Panama |
| *Paullinia glomerulosa* | Radlk. | Acevedo | 15194 | US | 3691466 | Panama |
| *Paullinia glomerulosa* | Radlk. | Chery | 20 | PMA | - | Panama |
| *Paullinia hystrix* | Radlk. | Acevedo | 14408 | US | 3620767 | Peru |
| *Paullinia hystrix* | Radlk. | Acevedo | 14417 | US | 3620775 | Peru |
| *Paullinia hystrix* | Radlk. | Acevedo | 14895 | US | 3680198 | Brazil |
| *Paullinia hystrix* | Radlk. | Acevedo | 14898 | US | 3680197 | Brazil |
| *Paullinia imberbis* | Radlk. | Acevedo | 14826 | US | 3680269 | Brazil |
| *Paullinia ingifolia* | Rich. & Juss. | Acevedo | 16067 | US | - | French Guiana |
| *Paullinia ingifolia* | Rich. & Juss. | Acevedo | 15248 | US | 3668951 | Panama |
| *Paullinia ingifolia* | Rich. & Juss. | Acevedo | 14812 | US | - | Brazil |
| *Paullinia ingifolia* | Rich. & Juss. | Acevedo | 15282 | US | 3668969 | Panama |
| *Paullinia jamaicensis* | Macfad. | Acevedo | 12035 | US | 3590021 | Jamaica |
| *Paullinia josecuatrii* | Macbr. | Acevedo | 14291 | US | 3629595 | Peru |
| *Paullinia killipii* | Macbr. | Acevedo | 14308 | US | 3625007 | Peru |
| *Paullinia killipii* | Macbr. | Acevedo | 14323 | US | 3620718 | Peru |
| *Paullinia largifolia* | Radlk. | Acevedo | 14920 | US | - | Brazil |
| *Paullinia latifolia* | Benth. ex Radlk. | Acevedo | 6038 | US | 3676038 | Suriname |
| *Paullinia latifolia* | Benth. ex Radlk. | Acevedo | 5835 | US | 3526316 | Suriname |
| *Paullinia leiocarpa* | Griseb. | Jansen-Jacobs | 3924 | US | 3359942 | Guyana |
| *Paullinia mazanensis* | Macbr. | Acevedo | 14364 | US | 3630454 | Peru |
| *Paullinia neglecta* | Radlk. | Nee | 37116 | US | 3174683 | Bolivia |
| *Paullinia neglecta* | Radlk. | Beck | 23548 | US | 3476039 | Bolivia |
| *Paullinia obovata* | (Ruiz & Pavón) Pers. | Acevedo | 14441 | US | 3592867 | Peru |
| *Paullinia obovata* | (Ruiz & Pavón) Pers. | Acevedo | 14382 | US | 3630472 | Peru |
| *Paullinia obovata* | (Ruiz & Pavón) Pers. | Acevedo | 14918 | US | - | Brazil |
| *Paullinia olivacea* | Radlk. | Acevedo | 14390 | US | 3620745 | Peru |
| *Paullinia paullinoides* | Radlk. | Acevedo | 7519 | US | 3330097 | Ecuador |
| *Paullinia paullinoides* | Radlk. | Acevedo | 14860 | US | 3680190 | Brazil |
| *Paullinia pinnata* | L. | Chery | 41 | PMA | - | Panama |
| *Paullinia pinnata* | L. | Acevedo | 6679 | US | 3580781 | Bolivia |
| *Paullinia pinnata* | L. | Kayombo | 1024 | US | 3316618 | Tanzania |
| *Paullinia pinnata* | L. | Pace | 192 | US | 3677377 | Brazil |
| *Paullinia pseudota* | Radlk. | Acevedo | 3708 | US | 3212304 | Brazil |
| *Paullinia pseudota* | Radlk. | Perdiz | 808 | CEPEC | 128946 | Brazil |
| *Paullinia pseudota* | Radlk. | Roque | 2594 | ALCB | 93954 | Brazil |
| *Paullinia revoluta* | Radlk. | Perdiz | 487 | US | 3627818 | Brazil |
| *Paullinia revoluta* | Radlk. | Perdiz | 822 | CEPEC | 128960 | Brazil |
| *Paullinia rubiginosa* | Cambess. | Acevedo | 14366 | US | 3630452 | Peru |
| *Paullinia rubiginosa* | Cambess. | Perdiz | 699 | - | - |  |
| *Paullinia rubiginosa* Cambess. subsp*. setosa* | (Radlk.) Acev.-Rodr. | Acevedo | 11114 | US | 3569426 | French Guiana |
| *Paullinia rufescens* | Rich. ex Juss. | Acevedo | 11120 | US | 3569432 | French Guiana |
| *Paullinia rugosa* | Benth. ex Radlk. | Chery | 25 | PMA | - | Panama |
| *Paullinia rugosa* | Benth. ex Radlk. | Acevedo | 15179 | US | 3691473 | Panama |
| *Paullinia rugosa* | Benth. ex Radlk. | Acevedo | 14807 | US | 3680195 | Brazil |
| *Paullinia selenoptera* | Radlk. | Acevedo | 14813 | US | - | Brazil |
| *Paullinia serjaniifolia* | Tria. & Planch. | Acevedo | 15270 | US | 3668972 | Panama |
| *Paullinia simulans* | Macbr. | Acevedo | 15235 | US | 3668974 | Panama |
| *Paullinia* sp. |  | Breedlove | 72699 | UCBG | 92.0509 | Mexico |
| *Paullinia* sp. |  | Chery | 46 | PMA | - | Panama |
| *Paullinia* sp. |  | Acevedo | 16064 | US | - | French Guiana |
| *Paullinia* sp. |  | Acevedo | 16089 | US | - | French Guiana |
| *Paullinia* sp. |  | Acevedo | 15718 | US | - | Colombia |
| *Paullinia* sp. |  | Acevedo | 15706 | US | - | Colombia |
| *Paullinia* sp. |  | Acevedo | 12335 | US | 3526309 | French Guiana |
| *Paullinia* sp. |  | Acevedo | 11105 | US | 3569445 | French Guiana |
| *Paullinia sphaerocarpa* | Rich. & Juss. | Acevedo | 15247 | US | 3668952 | Panama |
| *Paullinia spicata* | Benth. | Acevedo | 12344 | US | 3526306 | French Guiana |
| *Paullinia sprucei* | Macbr. | Acevedo | 14922 | US | - | Brazil |
| *Paullinia stellata* | Radlk. | Torke | 615 | MO | 6709351 | Bolivia |
| *Paullinia stipitata* | Cuatre. | Acevedo | 15280 | US | 3668967 | Panama |
| *Paullinia stipitata* | Cuatre. | Acevedo | 15242 | US | 3668954 | Panama |
| *Paullinia subrotunda* | (Ruiz & Pavón) Pers. | Melo | 402 | US | 3682609 | Brazil |
| *Paullinia ternata* | Radlk. | Perdiz | 824 | CEPEC | 128962 | Brazil |
| *Paullinia tomentosa* | Jacq. | Acevedo | 12236 | US | - | Mexico |
| *Paullinia trigonia* | Vell. | Perdiz | 673 | CEPEC? | - | Brazil |
| *Paullinia turbacensis* | Kunth | Acevedo | 15176 | US | 3691475 | Panama |
| *Paullinia turbacensis* | Kunth | Chery | 10 | PMA | - | Panama |
| *Paullinia turbacensis* | Kunth | Chery | 13 | PMA | - | Panama |
| *Paullinia turbacensis* | Kunth | Chery | 24 | PMA | - | Panama |
| *Paullinia verecunda* | Standl. | Acevedo | 15256 | US | 3683512 | Panama |
| *Paullinia verecunda* | Standl. | Acevedo | 15234 | US | 3668959 | Panama |
| *Serjania atrolineata* | C.Wright | Acevedo | 15195 | US | 3691465 | Panama |
| *Serjania atrolineata* | C.Wright | Chery | 42 | PMA |  | Panama |
| *Serjania circumvallata* | Radlk. | Acevedo | 15203 | US | 3691463 | Panama |
| *Serjania cornigera* | Turcz. | Chery | 36 | PMA | 112573 | Panama |
| *Serjania deltoidea* | Radlk. | Acevedo | 15725 |  |  | Colombia |
| *Serjania fluminensis* | Acev.Rodr. | - | - | - | - | Brazil |
| *Serjania membranacea* | Splitg. | Acevedo | 15227 | US | 3691479 | Panama |
| *Serjania mexicana* | (L.) Willd. | Chery | 23 | PMA | - | Panama |
| *Serjania mexicana* | (L.) Willd. | Chery | 45 | PMA | - | Panama |
| *Serjania paucidentata* | DC. | Chery | 34 | PMA | - | Panama |
| *Serjania pyramidata* | Radlk. | Chery | 29 | PMA | 112578 | Panama |
| *Serjania reticulata* | Cambess. | Torke | 508 | US | 3677589 | Bolivia |
| *Serjania rhombea* | Radlk. | Acevedo | 15213 | US | 3691459 | Panama |
| *Serjania salzmanniana* | Schltdl | Perdiz | 806 | US | 3627810 | Brazil |
| *Serjania* sp. |  | Chery | 18 | PMA | - | Panama |
| *Serjania trachygona* | Radlk. | Acevedo | 15222 | US | 3668956 | Panama |
| *Serjania pyramidata* | Radlk. | Acevedo | 15243 | US | 3668953 | Panama |
| *Talisia nervosa* | Radlk. | Acevedo | 15190 | US | 3683511 | Panama |
| *Talisia* sp. |  | Perdiz | 939 | US | 3627815 | Brazil |
| *Thinouia mucronata* | Radlk. | Zardini | 13513 | US | 3239039 | Paraguay |
| *Urvillea ulmacea* | Kunth | Balick | 3349 | US | 3296216 | Belize |
