## Appendix A for "Phylogeny of *Paullinia* L. (Paullinieae: Sapindaceae), a diverse genus of lianas with rapid fruit evolution"

**Appendix A**
Locus characteristics, PCR amplification and sanger sequence primer validation results, and MiSeq sequencing success results.

| **Locus** | **Region** | **Primer Validation/ Sanger Sequenced Successfully** | **Illumina MiSeq Reads Attained** | **Locus Included in Final Alignment** | **%Pairwise Identity** | **Alignment Length** | **#OTUs** |
| --- | --- | --- | --- | --- | --- | --- | --- |
| orange1.1g002083m (intron9) | Nuclear Intron | + | + | + | 86.8 | 662 | 101 |
| orange1.1g015495m (intron8) | Nuclear Intron | + | + | + | 73.2 | 237 | 15 |
| orange1.1g027952m (intron5) | Nuclear Intron | + | + | + | 85.4 | 605 | 54 |
| orange1.1g022777m (intron3) | Nuclear Intron | + | + | + | 80.6 | 847 | 36 |
| orange1.1g016982m (intron11) | Nuclear Intron | + | + | + | 94.7 | 176 | 122 |
| orange1.1g009973m (intron5) | Nuclear Intron | + | + | + | 84.9 | 316 | 40 |
| orange1.1g030977m (intron1) | Nuclear Intron | + | + | + | 93.2 | 285 | 82 |
| orange1.1g036770m (intron27) | Nuclear Intron | + | + | + | 81.1 | 703 | 88 |
| orange1.1g019384m (intron3) | Nuclear Intron | + | + | + | 78.8 | 693 | 35 |
| psbA-trnH | chloroplast | N/A | + | + | 89.6 | 594 | 119 |
| ITS | nrDNA | + | N/A | + | 84.9 | 763 | 122 |
| orange1.1g028997m (intron4) | Nuclear Intron | + | + |  |  |  |  |
| orange1.1g016982m (intron11) | Nuclear Intron | + | + |  |  |  |  |
| orange1.1g022600m | Nuclear Exon | + |  |  |  |  |  |
| orange1.1g011087m | Nuclear Exon | + | + |  |  |  |  |
| orange1.1g039733m | Nuclear Exon | + |  |  |  |  |  |
| orange1.1g001405m | Nuclear Exon | + | + |  |  |  |  |
| orange1.1g028318m | Nuclear Exon | + | + |  |  |  |  |
| orange1.1g004101m | Nuclear Exon | + |  |  |  |  |  |
| orange1.1g008050m | Nuclear Exon | + | + |  |  |  |  |
| orange1.1g000832m | Nuclear Exon | + |  |  |  |  |  |
| orange1.1g004904m | Nuclear Exon | + | + |  |  |  |  |
| orange1.1g013475m | Nuclear Exon | + |  |  |  |  |  |
| orange1.1g022288m | Nuclear Exon | + |  |  |  |  |  |
| orange1.1g000428m | Nuclear Exon | + | + |  |  |  |  |
| orange1.1g011916m | Nuclear Exon | + |  |  |  |  |  |
| *rpo*B | chloroplast | N/A |  |  |  |  |  |
| trnD-T | chloroplast | N/A |  |  |  |  |  |
| trnL-F | chloroplast | N/A |  |  |  |  |  |

Forward and Reverse primers of loci that successfully amplified by PCR + sanger sequenced.

| **Locus** | **Forward Primer** | **Reverse Primer** | **Citation** |
| --- | --- | --- | --- |
| orange1.1g002083m (intron9) | CATATGCAGTTACAGCACTAATGA | AATCTCAACAGCATGAGCATC | Chery et al. (2017) |
| orange1.1g015495m (intron8) | CTGCTGGAAATGCCTCTAGC | CTGAGCAGCGTCAGCATATC | Chery et al. (2017) |
| orange1.1g027952m (intron5) | TGGTTTTGATTGATGCAAGTG | GCATCTTCCCACCAAGGATA | Chery et al. (2017) |
| orange1.1g022777m (intron3) | GGAGGATTTCAATGAGGCTCT | TCTCAGCATAATCAGACCTGTG | Chery et al. (2017) |
| orange1.1g016982m (intron11) | CATTCCGTGATTTGCCTCTT | TCCATATTCCTGTTTCATCTGC | Chery et al. (2017) |
| orange1.1g009973m (intron5) | AGTGGAACTGCTTCGCAAGT | TGCATATGGGTTATAGCCTTGA | Chery et al. (2017) |
| orange1.1g030977m (intron1) | ACCGCCTCCCTATTACAGACTCTACA | TGGGTAAAGCTGACGCACTCCTTG | Chery et al. (2017) |
| orange1.1g036770m (intron27) | TGAAGCCATTTTCCAGTGCACATT | ACCGAATCAATGCAGGAAAACAGTGA | This study |
| orange1.1g019384m (intron3) | TGCATTCAAATGTCACCGAAAATCA | ACCATCACATCCTCCAGTAGCAAA | This study |
| psbA-trnH | CGCGCATGGTGGATTCACAATCC | GTTATGCATGAACGTAATGCTC | (Sang et al. 1997; Tate and Simpson 2003) |
| ITS | TCCTCCGCTTATTGATATGC | CCTTATCATTTAGAGGAAGGAG | (Whites et al. 1990; Stanford et al. 2000) |
| orange1.1g028997m (intron4) | AAAGAGTCCAAACCAACAATTC | TAAGCAGCACTTTTCCCACA | Chery et al. (2017) |
| orange1.1g016982m (intron11) | GACCAAATCATTTCTGGGATAGAC | TAGCCTAAGGATAACAAGGATGG | This study |
| orange1.1g022600m | TTCGTTGTCTATCAATTCTGAA | TTAGAAGCAGTGTGAGTAATTT | This study |
| orange1.1g011087m | ATGATGCAGGACATGATAAAG | TCTTTATAATTCCCATCATCTCTC | This study |
| orange1.1g039733m | GATTTAAAATGTGATTGTCAACAT | ATTTGCTAAAAACCCTAAAGTTTA | This study |
| orange1.1g001405m | CATGTACTGGATAGCATCTAC | CTTAAGTCCAAAACTCCTCTC | This study |
| orange1.1g028318m | TGTGATGATATTAACTTGGATTTT | AGGTCTATGATAAAATTATCCCAA | This study |
| orange1.1g004101m | TTACACAGAGTCAGAAATTTTTAC | TACTTCCAAGTTACTGAAATGG | This study |
| orange1.1g008050m | CTTGAGTGGAGTTGTTAAGATAT | CTTCTAATGCAAGAATACTTCTC | This study |
| orange1.1g000832m | ATTGCATTTACTTTTGGACTTG | ATTAAACTGGAAAGTTCGAATATC | This study |
| orange1.1g004904m | CGTAATGCTTTGTATAACATACA | CAACTTGTTCTTCTGACAAAG | This study |
| orange1.1g013475m | ATGGATCAATCAATCTGCATAT | ATAGAATTCTCACTCATATTGACT | This study |
| orange1.1g022288m | CATCTCTTAGATCTGATATTGAGA | AGTTTTCAATTCTAGGTCAGTT | This study |
